# Injectable adhesive hydrogel-drug complexes synchronize the release of chemoimmunotherapy to treat brain tumors in mice

**DOI:** 10.1101/2025.03.31.646388

**Authors:** Michelle Z. Dion, Alexander M. Cryer, Daniel Dahis, Pere Dosta, Andrea Li, Maria Poley, Nuria Puigmal, Ivana Ling, Sofia Carrasco, Alexa Hinojosa, Yael Soria, Betty Tyler, Henry Brem, Natalie Artzi

## Abstract

Brain tumor therapy remains limited by an immunosuppressive, monotherapy-resistant tumor microenvironment and by a paucity of technologies capable of controlling therapy delivery kinetics to the tumor. Here, we describe tissue-adhesive hydrogel-drug complexes (HDCs) for the controlled release of combination therapies synchronized with immune cycle responses following intracranial administration. Injected adhesive HDCs enabled the controlled co-delivery of multiple payloads (chemotherapy, stimulator of interferon genes agonist cyclic dinucleotide nanoparticles, immune checkpoint blockade antibodies), leading to enhanced long-term survival (>80%) and protection from contralateral hemisphere rechallenge after a single dose in multiple syngeneic orthotopic glioblastoma mouse models. Mechanistic studies revealed that tumor rejection is a consequence of reprogramming the tumor microenvironment, driven by three key phases: the initial rapid expression of inflammatory cytokines, notably IFN-γ, and tumor sensitization with antigen exposure during chemoimmunotherapy release, followed by the recruitment and subsequent sustained activation of antigen-presenting and effector cells facilitated by the hydrogel’s multiweek delivery period. These findings point to tissue-adhesive HDCs, and therapeutic release synchronized with biological responses as key considerations for realizing robust immune activation without provoking toxicity in the treatment of solid tumors.

## Introduction

Glioblastoma (GBM) is an aggressive and uniformly lethal form of brain cancer that carries a dismal prognosis despite multimodal therapy, with less than 10% of patients surviving for more than five years^1^. Recent advances in cancer therapy, such as targeted therapies and immunotherapies, have also failed to significantly extend patient survival, except for tumor-treating fields^2,3^. The development of more effective GBM therapies is limited by ineffective drug delivery to brain tumors and an immunosuppressive, heterogeneous tumor microenvironment (TME). Drug delivery to the brain is complicated by the presence of the blood-brain barrier (BBB), which prevents the effective penetration and retention of systemically administered therapies in the tumor^4^, limiting the treatment modalities available to patients. Unique anatomy also distinctly regulates the interaction between the central nervous system, including the brain parenchyma, and the peripheral immune system^5^, posing a significant challenge to the sequential coordination of immune biological processes across these compartments, which is critical for eliciting robust and durable anti-tumor immune responses^6^. Furthermore, even with effective drug delivery, there is a scarcity of drugs that can convert the hostile TME to an immunostimulatory state, given the abundance of inflammation-resolving myeloid cells^7^, the low effector T and natural killer (NK) cell infiltration in the TME^8^, and systemic immunosuppression, including iatrogenic effects^9^. The therapeutic window of these types of immunotherapies, such as STING agonists^10,11^, are narrow, particularly in the central nervous system where studies have implicated cGAS-STING signaling and type I IFN production in pathologic neuroinflammation^12^, neurodegeneration^13^, and peripheral tumor brain metastasis^14^, requiring an exquisite balance between protective immune activation and neurotoxic effects. These challenges emphasize the critical need for new technologies to treat GBM.

Local drug delivery can overcome these challenges by concentrating therapy at the tumor site, eliminating the need to cross the BBB and reducing the therapy’s distribution to off-target tissues, allowing effective therapeutic windows to be realized. This affords the opportunity to select therapies that can address GBM’s significant immunosuppression and intratumoral heterogeneity. The clinical feasibility and potential of local delivery approaches has been demonstrated by FDA-approved Glidael™, and recent Phase I engineered cell therapy trials^15–17^. While local polymer-based delivery strategies have been explored for the treatment of GBM^18–19^, numerous challenges remain including poor implant conformation to, and retention at, the pathologic site, difficult manufacturing and clinical administration procedures, and insufficient control over drug release kinetics, preventing the use of potent combination therapies.

Here, we develop tissue adhesive dextran-dendrimer hydrogel-drug complexes (HDCs) for the controlled delivery of an *in-situ* vaccine cocktail consisting of immunogenic cell death (ICD)-inducing chemotherapy doxorubicin (DOX), stimulator of interferon genes (STING) agonist cyclic dinucleotide nanoparticles (CDN-NPs), and immune checkpoint blockade (ICB) antibody, anti-Programmed Death-1 (aPD-1), to brain tumors following intracranial injection or spraying (**Fig. 4.1A**). Adhesive hydrogels can improve drug penetration of the tumor by effectively contouring to the tumor tissue to reduce the required drug diffusion distance and to maximally coat the target surface^20,21^. We leveraged the multivalency of hydrogel polymers to enable concurrent tissue adhesion, hydrogel cohesion, and drug interactions for controlled release without sacrificing hydrogel integrity. *In situ* vaccination, which combines a cytotoxic agent with an immunostimulatory agent, may be a powerful strategy by which to address the shortcomings of existing molecular therapies due to its ability to control tumor burden through ICD while concurrently reshaping, recruiting, and educating the immune system to eliminate residual disease. However, little is known about the impact of *in situ* vaccine release kinetics on the formation of immune responses and memory, particularly for brain tumors. We hypothesized that drug-hydrogel interactions could be modulated to allow for the delivery of immunomodulatory therapies in a manner that was synchronous with incited immune responses, enhancing the potency of the *in situ* therapeutic regimen. Our studies in orthotopic mouse models demonstrated that adhesive HDC *in situ* vaccines stimulate and maintain diverse immune responses that promote brain tumor control and elimination without inducing central nervous system autoimmunity. This approach overcomes drug delivery and immunobiological challenges for successful brain cancer therapy and demonstrates the use of HDCs to synchronously complement biological immune responses through controlled intracranial combination therapy delivery.

**Figure 1:**
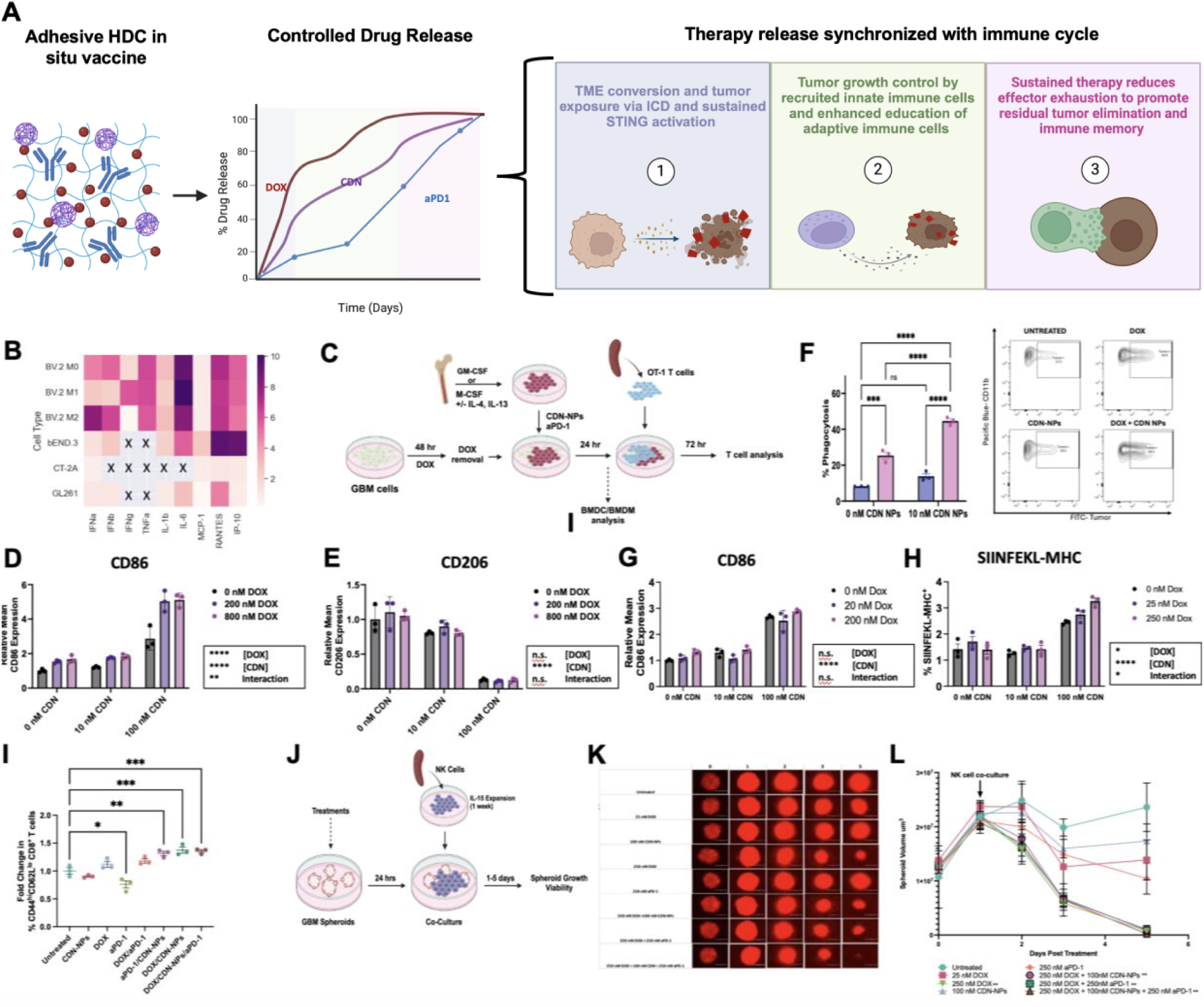
In situ vaccine payloads reshape myeloid polarization and tumor recognition by immune cells, leading to enhanced effector cell activity *in vitro*. **a**, Schematic illustration of HDC codelivery of multiple payloads for synchronous delivery with incited immune responses. **b,** Quantification of cytokines expressed by M0, M1 or M2-BV.2 microglia, bEND3 cells, GL261-luc2 cells, or CT-2A-luc cells following treatment with CDN-NPs using LegendPlex assay. Samples indicated with X were below the assay level of detection. Values are presented as log2-fold change relative to the average of untreated cells. **c**, Schematic of co-culture experiments used for evaluation of BMDM activation and phagocytosis (**d**-**f**), BMDC activation and antigen cross-presentation (**g**,h), and T cell proliferation and maturation (**i**). **d, e,** Flow quantification of the fold-change of M1-like (CD86, **e**) and M2-like (CD206, **f**) marker expression by M2-polarized BMDM cells in co-culture with tumor cells following treatment with chemoimmunotherapies. **f**, Flow quantification and representative flow plots of BMDM phagocytosis of GBM tumor cell lines following treatment with chemoimmunotherapies. **g**, **h**, Flow quantification of the relative expression of CD86 activation marker (**g**) and of SIINFEKL-MHC-I complex (**h**) on BMDCs following treatment in co-culture with tumor cells. **i**, Flow quantification of the fold change in effector memory CD8 T cells (CD8^+^CD44^hi^CD62L^lo^) following BMDC-tumor-T-cell co-culture (n=3). **j**, Schematic of primary NK cell co-culture with tumor spheroids to evaluate NK cell activity. **k**, Representative fluorescence images of GBM spheroids following treatment with and without NK cells. **l**, Spheroid volume following treatments over the course of five days with and without NK cells, calculated using ImageJ analysis. Data are presented as the mean± s.e.m. Statistical significance was determined using one-way or two-way ANOVA. *P<0.05, **P<0.01, ***P<0.001, ****P<0.0001.

### *In situ* vaccine reprograms myeloid cells and promotes MHC-dependent and MHC-independent anti-tumor immune responses *in vitro*

Therapy selection is critical to successful GBM therapy, however, the available therapeutics for selection are limited due to the paucity of therapies that cross the BBB and that have an attainable therapeutic window. This is further exacerbated for combination therapy approaches, which are likely necessary to address the cellular heterogeneity and multifaceted immunosuppression of GBM. We hypothesized that the timed release of combination therapy components to be synchronous with the immune responses they modulated would improve therapeutic outcomes (**Fig. 1A**). To explore this hypothesis, we began by assessing the effects of our proposed combination therapy *in vitro*.

We reasoned that the initial phase of therapy would need to achieve (1) effective TME reprogramming by repolarizing myeloid cells and driving immune cell recruitment through cytokine expression, and (2) control tumor growth and immune exposure through ICD. We previously developed CDN-NPs consisting of a potent CDN molecule conjugated to poly-beta-amino ester peptide-modified polymers via a cathepsin sensitive linker^22^, which facilitates CDN release from the endosome upon cellular uptake and improves NP drug loading and stability. We first showed that CDN-NP treatment could activate interferon regulatory factors in STING dual reporter RAW cells and induce inflammatory cytokine production by multiple cell types *in vitro*, with the highest fold-changes induced in immune cells and endothelial cells (**Fig. 1B, Fig. S1A- C**). We next demonstrated that DOX was a more potent chemotherapy than existing GBM chemotherapies (temozolomide and carmustine) in 2D and 3D *in vitro* GBM models (**Fig. S2A-C**) and that it also induced ICD of tumor cells (**Fig. S2D-E,G**). We observed that DOX treatment enhanced cell surface calreticulin expression and other immunoregulatory markers, including PD-L1 and PD-L2, in a dose-dependent and time-dependent fashion (**Fig. S2F-H**). We next explored the effects of chemoimmunotherapy in bone-marrow derived macrophages (BMDMs) in co-culture with tumor cells to understand if changes in intercellular communication and cell state in response to the therapy could impact tumor cell survival and immune activity (**Fig. 1C**). Tumor-associated myeloid cells are a significant portion of cells in the GBM TME that support numerous mechanisms of tumor growth through their immunosuppressive phenotype^7,23^. One strategy to promote anti-tumor immune responses in GBM is to eliminate or repolarize TAMs to a pro-inflammatory state. We therefore examined activation (CD86, CD80) and regulatory marker expression (CD206) by IL-4, IL-13 polarized BMDMs in co-cultures with DOX-treated tumor cells. We found that, like in monocultures of BMDMs and microglia (**Fig. S1D-E**), CDN-NP monotherapy was able to enhance expression of activation markers (CD86, CD80) and decrease expression of the regulatory marker CD206 in a dose-dependent fashion (**Fig. 1D-E**). DOX treatment enhanced the expression of activation markers by BMDMs, but did not have a significant effect on CD206 expression. Combination therapy resulted in significantly higher activation marker expression than either monotherapy alone. Based on the ability of combination culture to prime macrophage activation, we next assessed whether combination therapy could enhance macrophage effector function by quantifying changes in phagocytosis in tumor-BMDM co-cultures following treatments. DOX treatment of tumor cells increased their phagocytosis by BMDMs in a dose-dependent fashion (**Fig. 1F, S3A**). CDN-NP treatment had a more modest effect on BMDM phagocytosis of tumor cells but further enhanced phagocytic levels when combined with DOX treatment in tumor-BMDM co-cultures. Since microglia are the primary brain resident phagocytes and APCs, we also assessed phagocytic index in microglia-tumor cell co-cultures. DOX treatment enhanced microglia tumor cell phagocytosis, though CDN-NP therapy did not have a demonstrable effect on tumor phagocytosis alone or in combination in these cells (**Fig. S3B**). These data demonstrate that CDN-NP therapy can induce cytokine production as a monotherapy and myeloid reprogramming alone and in combination with DOX. We further show that combination therapy, driven by DOX, can increase macrophage phagocytosis of tumor cells in *in vitro* co-cultures.

We next assessed whether combination therapy could enhance the activation of bone-marrow derived DCs (BMDCs) and their ability to cross-present antigens when co-cultured with tumor cells. Like microglia and BMDMs, we found that CDN-NP and DOX monotherapies increased the maturation (as indicated by MHCII^+^CD86^+^) and activation (CD80^+^) of BMDCs in a dose-dependent fashion, however, distinctly, combination therapy did not result in further enhancement of maturation or activation (**Fig. 1G, Fig. S3C**). We also observed that combination therapy increased the phagocytosis of tumor cells by BMDCs (**Fig. S3D**). We next assessed whether this increase in maturation and activation of BMDCs was associated with increased tumor antigen cross-presentation when co-cultured with OVA-expressing tumor cells. BMDCs from co-cultures treated with combination therapy showed the highest level of antigen cross-presentation, followed by CDN-NP monotherapy (**Fig. 1H**). These results are consistent with the known effects of STING agonism on DCs^24^ and suggest that chemoimmunotherapy can improve the APC directed effects of CDN monotherapy. Antigen cross-presentation is required for the efficient priming of antigen-specific T cells. To determine if improved DC activation and function by combination therapy resulted in changes in T cell proliferation and maturation, we next isolated and added OT-I T cells to the BMDC-OVA-tumor cell co-cultures. Like for antigen cross presentation, combination therapies led to modest enhancements in the level of CD8 OT-1 T cell maturation as measured by the expression of effector memory markers **(Fig. 1I**).

While antigen-specific effector responses are an important component of anti-tumor immunotherapy, many tumors, including GBM, downregulate their antigen-presenting machinery^25^. Thus, when we were selecting an *in situ* vaccination regimen, we reasoned it would be important to stimulate multiple effector cell types with MHC-dependent and MHC-independent mechanisms. NK cells can mediate MHC-independent effector functions by inducing lysis of stressed cells^26^. Furthermore, STING agonism has been shown to promote anti-tumor NK cell responses in peripheral tumors^27–29^ and GBM^30^. Thus, we assessed the activation and cytotoxic activity of NK cells following treatment in co-culture with tumor spheroids (**Fig. 1J**). While treatment with therapies alone led to minimal changes in spheroid size (**Fig. S3E-F**), the inclusion of NK cells led to significant shrinkage of GBM spheroids treated with DOX-containing therapies relative to NK cell only treatment and treatment without NK cells (**Fig. 1K-L**). This was most significant in spheroids treated with the triple combination therapy, where the 3D structure was completely disintegrated by the treatment (**Fig. 1K**). Together, these data demonstrate that the combination therapy potently repolarizes myeloid cells and activates DCs to drive the expression of cytokines and chemokines for immune cell recruitment (**Fig. 1B**) and activation (**Fig. 1D,E,G**) and to cross-present tumor antigens (**Fig. 1H**). We show that treatment with immunogenic DOX sensitizes tumor cells to elimination by immune cells (**Fig. 1F, 1I, 1K**) activated by CDN-NP or recruited by chemokines expressed downstream of STING activation.

### Design and characterization of brain-adhesive injectable *in situ* vaccine HDCs

After confirming the activity of our combination therapy *in vitro*, we next tested its activity *in vivo* in the syngeneic orthotopic GL261 mouse model (**Fig. 2A**). While IT combination therapy significantly extended mouse survival, only one mouse survived tumor rechallenge (**Fig. 2A**). Thus, we hypothesized that controlling the release kinetics of the *in situ* vaccine therapy to align with biological responses to the therapy would improve the *in vivo* immune responses. To test this, we next engineered its release in injectable adhesive hydrogels *in vitro*. We used in vitro screening to identify a formulation with suitable gelation time, minimal swelling, tolerability, and prolonged degradation kinetics (over weeks) for intracranial use (**Fig. S4**). We elected to move forward with formulations displaying gelation times less than 1-minute post-mixing, that displayed minimal swelling, were well-tolerated, and that degraded over the course of several weeks in vitro (**Fig. S4**). We next characterized the physical and mechanical properties of the hydrogels when loaded with multiple therapies using cryo-scanning electron microscopy and rheometry (**Fig. 2B, Fig. S4**). These studies demonstrate that the hydrogels form gels with pores when loaded with the proposed therapies *in vitro* as expected. To ensure the applicability in the brain given its unique extracellular matrix, we tested the sprayability and adhesion of the hydrogel using ovine brain tissue under wet conditions. We have demonstrated that our hydrogel formulations are sprayable (**Fig. 2C**), which could facilitate their use as an adjuvant therapy to surgical resection in patients and maintain adhesion upon immediate immersion in a wet environment simulating surgical conditions (**Fig. S4G**). Finally, we characterized the release of the target therapeutic cargos from HDCs. We chemically modified DOX (aDOX) with aconitic anhydride to block its amine group from interacting with hydrogel polymers and modified the formulation pH to impact the polymer peptide amine protonation state. We demonstrated that the release profile of multiple payload types could be controlled by modulation of the drug’s or polymer’s accessible amine groups (**Fig. 2D-E**), which can be used to influence their release rate. Together, these studies demonstrate that our hydrogels gel upon spraying or injection, adhere to brain tissue to effectively coat complex tissue surfaces, and can control the release kinetics of diverse payloads.

**Figure 2:**
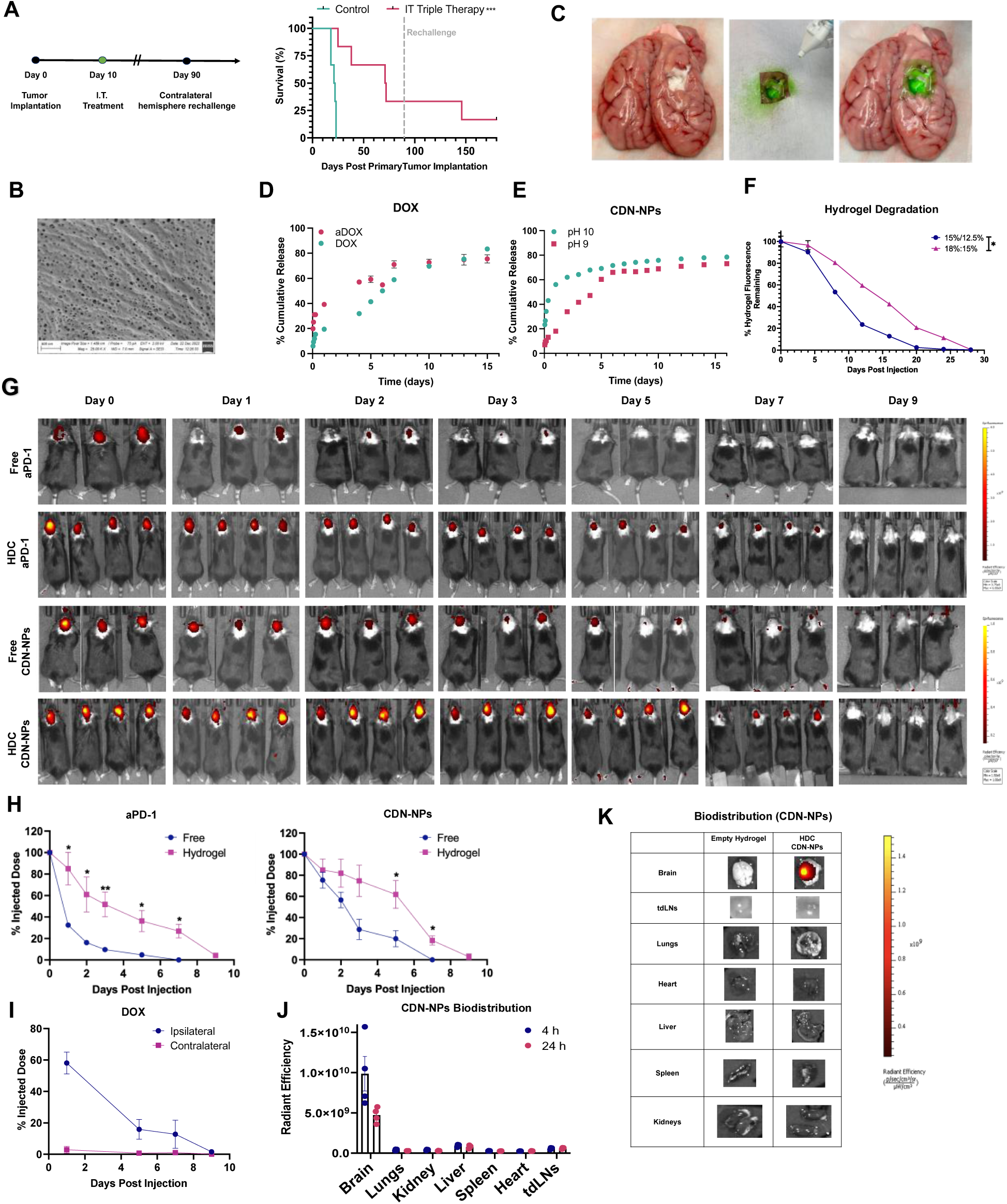
Characterization of injectable adhesive HDCs for the immune synchronous administration of in situ vaccine therapy in brain tumors. **a**, Kaplan-Meier survival curves of mice bearing GL261-luc2 orthotopic brain tumors following intracranial treatment upon primary challenge and contralateral rechallenge (n= 6 mice per group). **b**, Representative cryo-scanning electron microscopy images of the drug-loaded hydrogel formulation. **c**, Representative photographic images of fluoresceine-loaded hydrogels following spraying on wet porcine brain tissue ex vivo. **d, e,** Quantification of in vitro hydrogel release of DOX and amine-modified DOX (**d**) and CDN-NPs at different formulation pHs (**de** over time by fluorescence. **f**, Quantitative analysis of hydrogel degradation rate for fluorescently labeled hydrogels with different solid contents by fluorescent IVIS imaging over time in the TME of mice (n=4-6 mice per group). **g**, Representative IVIS images tracking the retention of fluorescently labeled antibodies or nanoparticles in the brains of mice following free therapy injection (IT) or hydrogel injection (n=3-4 mice per group). **h**, Quantification of the retention of fluorescently labeled therapies following intratumoral free or hydrogel injection by IVIS (n=3-4 mice per group). **i**, LC-MS/MS quantification of DOX in different hemispheres of the brain following hydrogel injection (n=3 mice per timepoint). **j**, Quantification of fluorescent CDN-NP distribution in organs 24 h post hydrogel treatment by ex vivo IVIS imaging (n=4 mice per treatment group). **k**, Representative images of organs at 24 h post hydrogel fluorescent CDN-NP treatment. Data are presented as the mean± s.e.m. Statistical significance was determined using the Student’s t-test. *P<0.05, **P<0.01, ***P<0.001, ****P<0.0001.

After confirming the performance of HDCs *in vitro*, we next assessed their drug delivery performance *in vivo*. First, we tracked the degradation of therapy-loaded fluorescently labeled HDCs in tumor-bearing mice using an In Vivo Imaging System (IVIS). Using this method, we confirmed that the fluorescent signal from the hydrogel persisted for up to three weeks and that hydrogel degradation is tunable to the formulation (**Fig. 2F**). To quantify the drug release kinetics of HDCs *in vivo*, we fluorescently labeled CDN-NPs and antibodies and injected the labeled materials intracranially in either an HDC formulation or in PBS and tracked their intracranial residence over time with IVIS. Formulating NPs or antibodies as HDCs led to significantly increased retention of therapeutic cargos over time in comparison to intracranially injection, even with the enhanced local cell uptake afforded by NP formulations (**Fig. 2G,H**). Next, we characterized the release of DOX from hydrogels using liquid chromatography tandem mass spectrometry (LC-MS/MS) following DOX extraction from the harvested brains. LC-MS/MS analysis showed that HDCs sustained delivery of DOX for over a week (**Fig. 2I**). Fluorescence microscopy imaging of the tumor at Days 2 and 5 post therapy also showed enhanced DOX retention in HDC-treated mice in comparison to mice treated with IT DOX (**Fig. S5B**). To understand if therapies distributed outside of the brain following HDC release, we further assessed therapy biodistribution (**Fig. 2I-K**). The amount of DOX was quantified in the ipsilateral hemisphere, contralateral hemisphere, and plasma. While there was quantifiable DOX present in the contralateral hemisphere (< 1% of the injected dose at peak on day 1), most of the dose remained in the ipsilateral hemisphere, and DOX concentration was below the level of quantification by LC-MS/MS in the plasma on each day assessed (days 1, 5, 7, 9) (**Fig. 2I**). We further quantified the biodistribution of CDN-NPs using IVIS fluorescence following hydrogel injection following mouse sacrifice and organ extraction at 4 h and 24 h post intracranial injection. Significant CDN-NP signal relative to background control mice was observed in the brain, but in no other major organs at either time point **(Fig. 2J, 2K**). This study confirms that, regardless of therapy type, HDC-mediated therapy remains localized to the brain, reducing the potential for adverse effects that may be seen with other administration methods. These results demonstrate that HDCs are biodegradable, biocompatible, and able to sustain therapy release *in vivo* following intracranial injection.

### HDC combination therapy eliminates established brain tumors in multiple syngeneic murine models and induces long-term protection from contralateral tumor rechallenge

After confirming that HDCs can sustain the release of target therapeutic cargos *in vivo* and that combination therapy enhances the activity of multiple target GBM immune populations *in vitro*, we performed efficacy studies in syngeneic, orthotopic GBM mouse models. Maximum tolerated dose studies in multiple models were used to determine the therapeutic doses (**Fig. S6**). For the GL261 model, HDC therapies or vehicle control were then administered via intracranial injection at the same location on day 10, following confirmation of tumor growth using IVIS imaging (**Fig. 3A**). In comparison to vehicle control (PBS or empty hydrogel), all HDC monotherapies and combination DOX/aPD-1 therapy delayed tumor growth as evidenced by IVIS bioluminescence, while all other HDC combination therapies lead to tumor regression, which was sustained in most mice in only the triple combination therapy (**Fig. 3B-C, S6**). These changes in tumor growth kinetics lead to significantly improved median overall survival in all HDC treatment groups relative to vehicle control (16 days, 0/14), and long-term survival in each HDC treatment group, with the highest median overall survival and percentage and number of long-term survivors seen in the triple therapy group (undefined m.s., 80%, 8/10) (**Fig. 3D**). Of the dual combination therapies, HDC DOX/CDN-NPs led to the longest median survival in comparison to CDN-NPs/aPD-1 and DOX/aPD-1 (68.5 days vs. 44.5 days vs. 27.5 days), though all dual combination therapies had similar long-term survivor rates (40% vs. 40% vs. 30%) (**Fig. 3D**). Transient weight loss was observed in mice receiving intracranial CDN-NPs, consistent with previous reports for STING agonist treatments^22^ (**Fig. S7**). These data support the efficacy and safety of hydrogel therapies. We next assessed whether long-term survivors developed protective memory responses by rechallenging long-term surviving mice with contralateral hemisphere tumors 90 days after treatment. All HDC treatment groups, except HDC aPD-1, had at least one mouse that was able to reject tumor upon rechallenge, with the best performance seen for treatments containing CDN-NPs where 100% of mice (HDC CDN-NPs or aPD-1/CDN-NPs or DOX/CDN-NPs) or 87.5% (7/8, triple therapy) of mice rejected tumors upon rechallenge (**Fig. 3E**). To understand how therapy release kinetics impacted therapeutic efficacy, we compared the efficacy of HDC and intratumoral therapies in the GL261 model. We found that HDC triple therapy improved median survival and overall survival in comparison to intratumoral triple therapy upon primary challenge (**Fig. 3E**). Mice treated with HDC triple therapy rather than intratumoral triple therapy also had higher rates of tumor rejection upon rechallenge (87.5%, 7/8) than intratumoral triple therapy (50%, 1/2), leading to a long-term survival of 70% of hydrogel treated mice in comparison to 16.7% of intratumoral treated mice (**Fig. 3F**). These data demonstrate that extended therapy release kinetics and the proposed combinations can extend the median survival of mice and improve long-term survival in a moderately immunogenic orthotopic GBM model.

**Figure 3:**
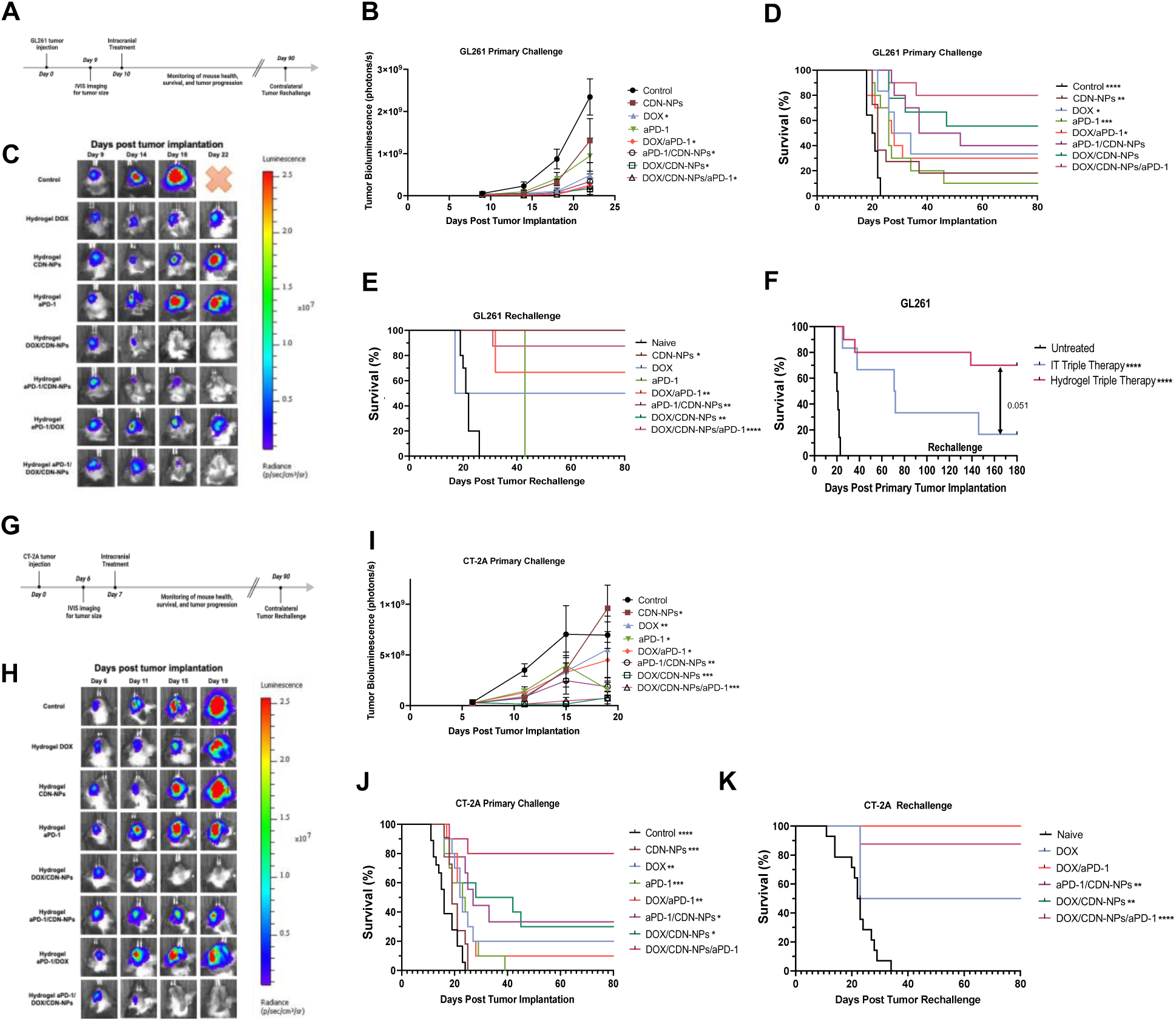
Anti-tumor efficacy of adhesive injectable HDCs co-delivering in situ vaccine payloads in mice with orthotopic brain tumors. **a**, Scheme of survival and rechallenge experiments evaluating the efficacy of hydrogel formulations in the GL261-luc2 model (1.5×10^5^ cells intracranially). **b,** Quantification of tumor bioluminescence over time following GL261-luc2 implantation by IVIS. **c**, Representative IVIS images depicting tumor bioluminescence in the GL261-luc2 model. **d**, Kaplan-Meier survival curves of mice bearing GL261-luc2 orthotopic brain tumors following intracranial hydrogel treatment upon primary challenge (n= 6-14 mice per group). **e**, Kaplan-Meier survival curves of long-term surviving mice (>90 days) following contralateral hemisphere rechallenge with GL261-luc2 brain tumors. **f**, Kaplan-Meier survival curves of mice bearing GL261-luc2 orthotopic brain tumors following intratumoral free drug and hydrogel triple therapy upon primary challenge and rechallenge, (n=6-10 mice per group). **g**, Scheme of survival and rechallenge experiments evaluating the efficacy of hydrogel formulations in the CT2A-luc model (1.5×10^5^ cells intracranially). **h**, Representative IVIS images depicting tumor bioluminescence in the CT2A-luc model. **i,** Quantification of tumor bioluminescence over time following CT-2A-luc implantation by IVIS. **j**, Kaplan-Meier survival curves of mice bearing CT2A-luc orthotopic brain tumors following intracranial hydrogel treatment upon primary tumor challenge (n= 10-18 mice per group). **k**, Kaplan-Meier survival curves of long-term surviving mice (>90 days) following contralateral hemisphere rechallenge with CT2A-luc orthotopic brain tumors in comparison to age-matched naïve mice. **b**,**i**, Data are presented as the mean± s.e.m. **d**,**e**,**f**,**j**,**k**, Statistical significance was determined relative to the control/naïve (**e**, **f**, **k**) or triple therapy **(d**, **i)** by the log-rank Mantel-Cox test for survival curves or by two-way ANOVA with multiple comparisons for tumor growth curves (**b**,**i**). *P<0.05, **P<0.01, ***P<0.001, ****P<0.0001.

After observing increased survival following HDC therapy in the GL261 model, we further characterized the therapeutic efficacy of our treatments in the more chemo-resistant, immunologically cold CT-2A GBM model^31^ (**Fig. 3G**). Delayed tumor growth was observed in HDC monotherapies and dual therapies in comparison to control mice, but complete tumor regression was not achieved in most mice treated with these therapies (**Fig. 3H-I**). Of the HDC dual therapies, HDC CDN-NPs/DOX was able to induce tumor regression transiently, but not able to control tumor burden in the long-term, resulting in extended median survival relative to other dual combination therapies (35 d vs. 27 d vs. 23 d), but not increased long-term survival (30% vs. 33.3% vs. 10%) (**Fig. 3J**). The combination of HDC aPD-1 and CDN-NPs led to significantly improved survival compared to HDC monotherapy of CDN-NPs or aPD-1. HDC triple therapy eliminated established CT-2A tumors in most mice, leading to significantly improved median and overall survival relative to all other groups studied (**Fig. 3J-K**). We also performed rechallenge studies for long-term survivors and found that HDC therapies prevented tumor regrowth in most mice with 13 out of 15 mice surviving long-term across all HDC groups, in comparison to no mice surviving tumor challenge in the naïve group (**Fig. 3K**). Together, these results show that HDC combination therapies are effective in eliminating established tumors and inducing immune memory in multiple orthotopic, syngeneic mouse models of GBM.

### HDC combination therapy increases myeloid activation and reprograms the TME in a host STING-dependent manner

To understand the mechanisms associated with HDC combination anti-tumor therapeutic efficacy, we first examined the contribution of host STING signaling by assessing the efficacy of HDC triple therapy in wild-type and *goldenticket* STING knockout (STING*^Gt^*) mice bearing CT-2A tumors (**Fig. 4A**). As before, treatment with HDC triple therapy led to tumor regression and increased long-term survival in wild-type mice in comparison to vehicle-treated mice (**Fig. 4B, 4C**). However, in CT-2A tumor-bearing STING*^Gt^* mice, combination therapeutic efficacy was significantly abrogated, with no significant differences in tumor growth or survival observed in comparison to mice treated with empty hydrogel vehicle control (**Fig 4B, 4C**). These results confirm that the activity of host cell STING signaling significantly contributes to combination therapy efficacy.

**Figure 4:**
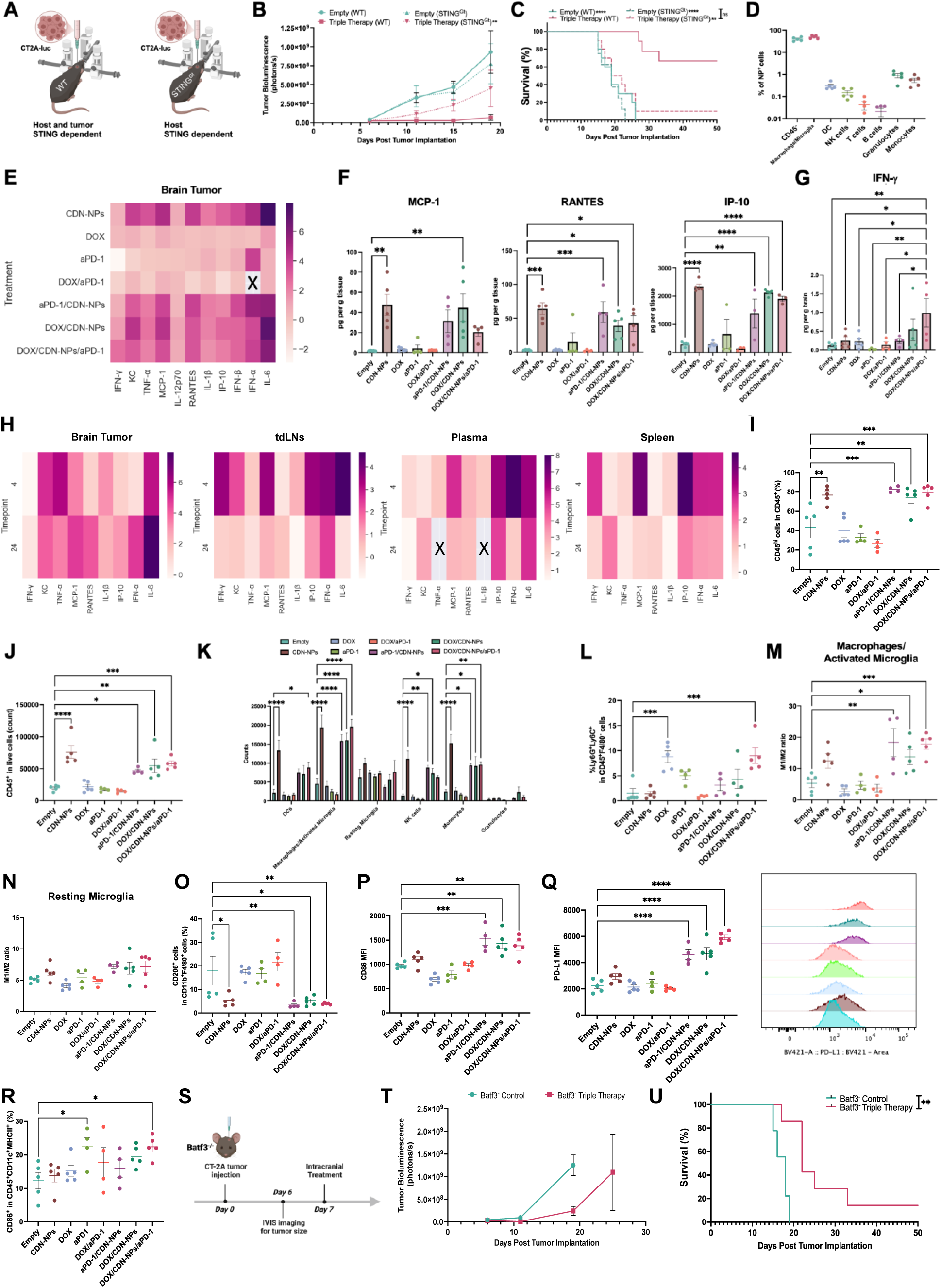
Adhesive HDCs release payloads that induce inflammatory cytokine production and activate antigen-presenting cells to potently remodel the TME through stimulation of host STING signaling. **a**, Schematic for determining the dependency of HDC triple therapy efficacy on host STING signaling. **b**, Quantification of tumor bioluminescence over time following CT2A-luc implantation (1.5×105 intracranially) by IVIS. **c,** Kaplan-Meier survival curves of wildtype (WT, solid lines) or goldenticket (STING^Gt^, dashed lines) mice bearing CT2A-luc orthotopic brain tumors following intracranial hydrogel treatment upon primary challenge (n= 8-10 mice per group). **d**, Cellular uptake of fluorescent CDN-NPs in the tumor. **e**, Cytokine profile in the CT2A tumor 24 h post treatment with different hydrogel formulations. Values are presented as log_2_-fold compared to CT2A tumor-bearing animals treated with empty hydrogels (n=4-5 mice per group). **f**,**g**, Quantifications of cytokines in the CT2A tumor per g tissue for each treatment using LegenedPlex assay (n=3-5 mice per group). **h**, Cytokine profile in different tissues 4 and 24 h post treatment with hydrogel CDN-NPs. Values are presented as log_2_-fold compared to CT2A tumor-bearing animals treated with empty hydrogels (n=4-5 mice per group). **i**, Flow quantification of CD45^+^ cells in live cells in the treated CT2A-tumor-bearing hemispheres of mice three days post hydrogel treatment (n=4-5 mice per group).j, Flow quantification of CD45^hi^ expression in CD45^+^ live cells in the tumor three days after hydrogel treatment (n=4-5 mice per group). **k**,**l**, Flow quantification of immune cell subsets within CD45^+^ live cells, including percentage of granulocytes (**l**), in tumors three days post hydrogel treatment (n=4-5 mice per group). **m**,**n**, Relative quantification of the ratio of M1-polarized (CD86^hi^CD80^hi^) to M2-polarized (CD206^hi^) macrophages/activated microglia (CD45^hi^ live cells) (**m**) and resting microglia (CD45^int^ live cells) (**n**) in the tumor three days after hydrogel treatment (n=4-5 mice per group). **o**, **p**, **q**, Flow quantification of M2 (CD206), M1 (CD86), and immunoregulatory (PD-L1) markers in macrophages/activated microglia (**o**) and resting microglia (**p**, **q**) in the tumor three days after hydrogel treatment (n=4-5 mice per group). **r**, Representative flow quantification of activated dendritic cells (CD45^hi^CD11c^+^MHCII^+^CD86^hi^) in the tumor three days after hydrogel treatment (n=4-5 mice per group). **s**, Schematic for the treatment of Batf3^-/-^ mice implanted with orthotopic CT-2A tumors. **t**, Quantification of tumor bioluminescence over time following CT2A-luc implantation (1.5×105 intracranially) by IVIS. **u,** Kaplan-Meier survival curves of Batf3^-/-^ mice bearing CT2A-luc orthotopic brain tumors following intracranial hydrogel treatment upon primary challenge (n= 7-9 mice per group). **a**,**e-q**, Data are presented as the mean± s.e.m. Statistical significance was determined by one-way ANOVA in comparison to empty hydrogel control mice unless otherwise indicated. **d**,**g**, Boxes containing x values are indicative of samples below the level of detection of the assay. **b,r**, Statistical significance was determined by the log-rank Mantel-Cox test relative to triple therapy hydrogel in wildtype mice or between groups as indicated. *P<0.05, **P<0.01, ***P<0.001, ****P<0.0001.

Given the impact of STING signaling on therapeutic efficacy, we next sought to determine the cellular distribution of CDN-NPs and the associated downstream spatiotemporal expression of pro-inflammatory cytokines following HDC combination therapy. Mice bearing CT-2A tumors were intracranially treated with HDC combination therapy containing fluorescently labeled CDN-NPs. After 24 hours, brain tumors were isolated from mice following transcardial perfusion, dissociated into single cell suspensions, and stained for analysis by spectral flow cytometry. CDN-NPs were principally internalized by CD45^-^ cells (39.9±3.6%) and tumor-associated macrophages and microglia (49.8± 3.3%), with less than 1% uptake seen in all other cell types analyzed in the brain TME (**Fig. 4.4D**). Similar cellular distributions were observed in the more immunogenic GL261 model (**Fig. S8**). As STING agonists are known to drive potent expression of anti-viral proinflammatory cytokines, we next characterized changes in pro-inflammatory cytokine expression in the TME in CT-2A tumor-bearing mice. In the TME, CDN-NP-containing HDCs increased expression of anti-viral cytokines, with highest fold-change in expression seen in IL-6 (**Fig. 4E**). IP-10, RANTES, and MCP-1, chemokines involved in the recruitment of monocytes, NK cells, T cells, DCs, and eosinophils, showed both high levels of expression and significant changes in expression relative to untreated controls in the TME following treatments (**Fig. 4F**). Increased type I interferons were also observed for all HDC CDN-NP-treated mice, as expected (**Fig. S9**). Though the change in expression was modest, increased IFN-γ expression was observed for HDC triple therapy treated mice relative to empty hydrogel and HDC CDN-NPs treated mice (**Fig. 4G**). To determine if cytokine expression profiles were spatiotemporally altered following local HDC triple therapy, cytokine profiles were also evaluated in the tumor, tdLNs, blood and spleen of CT-2A tumor-bearing mice to assess locoregional and systemic alterations at 4 h and 24 h post HDC triple therapy treatment (**Fig. 4H**). While peripheral immune organs and plasma did show increases in chemokine expression acutely at 4 h post HDC treatment which reflect the signature observed in the brain tumor, the expression was transient in these organs, whereas cytokine expression in the brain tumor continued to increase relative to empty hydrogel treated mice 24 h post treatment (**Fig. 4H**). This initial change could reflect the acute loss of CDN-NPs from burst release or their induced cytokines from the leaky tumor vasculature or alterations in intracranial fluid dynamics in response to surgical treatment. The 24 h timepoint is consistent with the biodistribution profile observed at 24 h, with the highest retention and distribution of NPs from the HDC seen in the brain (**Fig. 2J-K**).

To determine if CDN-NP uptake and combination therapy-induced cytokines impacted myeloid cell infiltration and phenotype, we next characterized changes in the frequency of tumor-associated myeloid cell populations and their expression of polarization markers following treatment. CT-2A tumors were treated with HDC therapies as previously described and then isolated and analyzed by spectral flow cytometry 72 h following treatment. All treatments containing CDN-NPs had an increase in CD45^+^ immune cell frequency (**Fig. 4I**), with a higher percentage of CD45^hi^ immune cells (indicative of peripherally derived myeloid and activated microglia) relative to CD45^int^ (indicative of resting microglia) immune cells (**Fig. 4J**). The increase in CD45^+^ immune cells was primarily driven by increases in DCs (CD45^+^CD11c^+^MHCII^+^), macrophages/activated microglia (CD45^hi^CD11b^+^F4/80^+^), NK cells (CD45^+^CD11b^-^NK1.1^+^), and monocytes (CD45^+^CD11b^+^Ly6C^+^Ly6G^+^) **(Fig. 4K**). Furthermore, elevated levels of granulocytes (CD45^+^CD11b^+^Ly6G^+^Ly6C^+^) were observed in DOX (8.8±1.2%) and triple therapy (9.0±1.6%) HDC-treated groups relative to empty hydrogel (1.5±0.9%) (**Fig. 4L**). Given the plasticity of macrophage and microglia in the TME^7,23,32^ and the observed impact of combination therapy on macrophage and microglia polarization *in vitro* (**Fig. 1D-E**, **S1, S3**), we further characterized the polarization of myeloid populations in the brain tumor by measuring expression of CD86 (pro-inflammatory, M1-like) and CD206 (M2-like, pro-healing) and deriving an M1/M2 ratio from these data (**Fig. 4M-N**). Significant changes to the M1/M2 ratio were only seen in macrophages and activated microglia in CDN-NPs containing combination therapy groups (**Fig. 4M**), largely driven by reduced CD206 expression (**Fig. 4O**). Slight increases in resting microglia M1/M2 ratio were observed, reflecting increases in CD86 expression by these cell types (**Fig. 4N, 4P**). Next, to understand how therapy impacted the expression of immunoregulatory markers, we measured the expression of PD-L1 across myeloid and stromal populations in the tumor. While macrophages and activated microglia expressed the highest levels of PD-L1 overall across all groups and cell types (**Fig. S10**), significantly higher expression of PD-L1 was only observed in HDC CDN-NP-containing combination therapies relative to empty hydrogel and HDC CDN-NPs in resting microglia (**Fig. 4Q**) and in granulocytes following HDC aPD-1/CDN-NPs therapy only (**Fig. S10**). These data demonstrate that HDC-mediated CDN-NP combination therapies increase myeloid activation in the TME, that is regulated, in part, by increased expression of PD-L1 by these cells, providing support for the use of local sustained ICB therapy targeting the PD-1/PD-L1 axis.

### DC recruitment and activation are induced by HDC chemoimmunotherapy efficacy, and their depletion abrogates therapy efficacy *in vivo*

Furthermore, within the myeloid compartment, DCs, particularly professional antigen cross-presenting conventional type 1 DCs (cDC1s), are known to contribute significantly to the activity of STING agonist therapies in peripheral tumors^24^; however, they are sparse in immunosuppressive tumors like GBM^7,31^. To examine the role of DCs in therapy efficacy, we characterized the frequency and activation of DC populations in the tumor and tdLNs. The frequency of DC (CD45^+^CD11c^+^MHCII^+^) was increased across HDC treatment groups containing CDN-NPs relative to empty hydrogel, though the overall percentage was decreased due to a significant influx of CD45^+^ cells within the TME in response to the therapy (**Fig. 4K**). Significantly increased activation of DCs (as indicated by CD86 expression) was only observed in HDC triple therapy and HDC aPD-1 groups (**Fig. 4R**), though all combination therapies showed increased levels of CD86 expression. We further characterized the tdLNs and spleen to determine if changes in DC or myeloid cell activity occurred in peripheral immune locations in response to the therapy. Significant differences in DC populations were not observed at these locations (**Fig. S11**). Given the minimal changes to DC activity observed by flow cytometry characterization, despite their known importance in peripheral tumors, we decided to further examine the effects of cDC1 activity on therapeutic efficacy using Batf3^-/-^ mice. We characterized the tumor growth and survival of Batf3^-/-^ CT-2A tumor-bearing mice treated with either empty hydrogel therapy or HDC triple therapy (**Fig. 4S**). Despite the limited alterations in DC frequency in the tdLNs, while triple therapy treatment was able to slow tumor growth and extend median survival relative to empty hydrogel controls (**Fig. 4T-U**), its therapeutic effects were significantly abrogated relative to triple therapy activity in wild-type mice. This suggests that cDC1 cells still contribute to the activity of HDC CDN-NP-containing combination therapies, perhaps at other timepoints or other locations of antigen presentation, such as local tertiary-lymphoid structures, the meninges^5^ or dural sinuses^33^, then those observed in the immunophenotyping studies, or through alternative mechanisms than those examined, such as through positive feedback loops with recruited NK cells^34^.

### HDC combination therapy increases infiltration and activation of NK cells, reduces T cell exhaustion, and is dependent on both cell types for efficacy

To understand if alterations in the TME affected immune effector cell recruitment and activation, we next assessed changes in the frequency and activation state of NK and T cell populations three- and eight-days post treatment in CT-2A-bearing mice (**Fig. 5A**). On day 3 post therapy, the frequency of NK cells and the percentage of NK cells (defined as NK1.1^+^ cells in CD45^+^CD11b^-^ cells) increased in the brain tumors of mice treated with HDC CDN-NPs monotherapy (14.8±2.3%) and combination therapies containing CDN-NPs (19.1±1.3%, 13.6±1.4%, 12.6±0.8%) (**Fig. 5B**). NK cells were also increased in the spleen, but not the tdLNs, following HDC triple therapy on day 3 relative to empty hydrogel-treated mice (**Fig. S4.11**), which complements the acute changes in cytokine profiles observed in peripheral immune organs (**Fig. 4H**). Significantly decreased PD-L1 expression was only observed in NK cells from mice treated with HDC DOX/CDN-NPs (**Fig. S11**). The percentage of NK cells in GBM immune infiltrates was also increased at day 8 post therapy (though to a lesser extent) for HDC CDN-NPs treated mice (8.0±1.1%) relative to empty hydrogel-treated mice (4.3±0.8%) (**Fig. 5B**). While none of the combination therapies sustained significantly increased levels of NK cells relative to mice treated with empty hydrogel control on day 8, increased NK cell activation (as indicated by expression of CD69) was observed in HDC CDN-NPs (54.7±3.3%), HDC DOX/CDN-NPs (59.0±3.6%), and HDC triple therapy (55.6±1.8%) (**Fig. 5C**). There were no significant changes to NK cell expression of PD-1 following therapy (**Fig. S11**). While immune profiling by flow did not show significant differences in NK cell recruitment and activation between hydrogel CDN monotherapy and combination therapies, we hypothesize that the release of DOX could induce both ICD and injury in tumor cells that survive. This low-dose injury phase may result in higher expression of cell stress molecules, such as calreticulin (**Fig. S2**), that can improve the identification and clearance of residual tumor cells by MHC-independent cytotoxicity as we observed *in vitro* (**Fig. 1K-L**).

**Figure 5:**
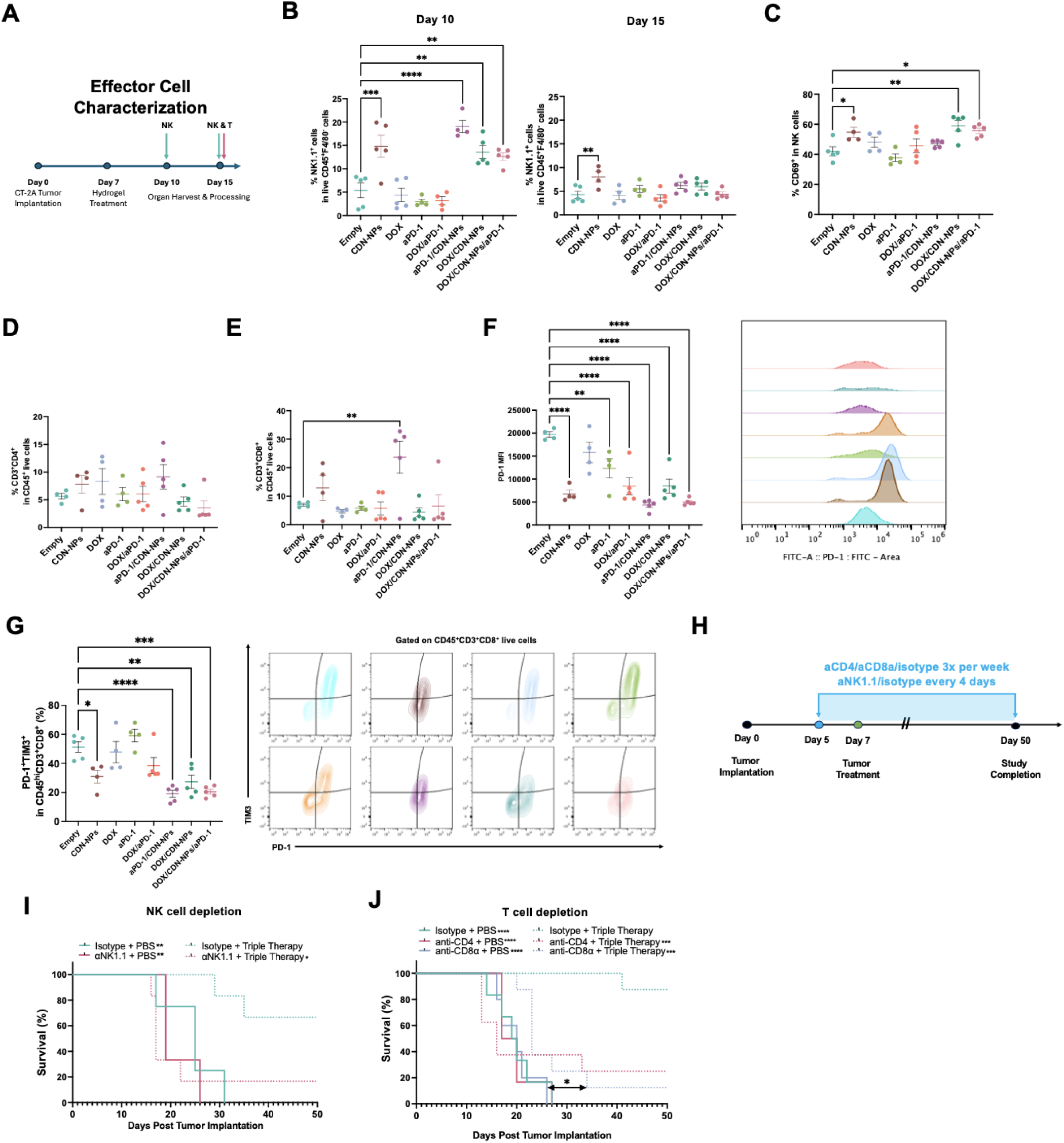
Sustained release of in situ vaccination regimen enhances activation and reduces exhaustion of NK and T effector cells. **a**, Schematic for flow cytometry experimental analysis of NK and T effector cells three days and eight days post hydrogel treatment in an orthotopic CT2A-luc model. **b**, Flow quantification of the percentage of NK1.1^+^ cells in CD45^hi^F4/80^-^ live cells on days three and eight post treatment. **c**, Representative flow quantification of early activation in NK cells as assessed by CD69 expression eight days post treatment. **d**, **e**, Percentage of CD4 (CD3^+^CD4^+^) and CD8 (CD3^+^CD8^+^) T cells in the tumor eight days following treatment by flow cytometry. **f-g**, Quantification of PD-1 and TIM-3 exhaustion marker expression on CD8 T cells in the tumor and the associated representative flow plots as assessed by flow cytometry. **h**, **i**, Kaplan-Meier survival curves of mice bearing CT2A-luc orthotopic brain tumors following intracranial hydrogel treatment upon primary challenge with antibody-mediated depletion of NK cells (**h**) and CD4 and CD8 T cells **(i**) (n= 5-8 mice per group). **b-h**, Data are presented as the mean± s.e.m. Statistical significance was determined by one-way ANOVA in comparison to empty hydrogel control mice unless otherwise indicated. **h**,**i**, Statistical significance was determined by the log-rank Mantel-Cox test between groups as indicated. *P<0.05, **P<0.01, ***P<0.001, ****P<0.0001.

Therapy-induced alterations to the frequency and state of T cell populations in the brain, tdLNs, and spleen were also characterized on day 8. Despite the high levels of CXCL10 induced by STING-containing therapies, the percentage of CD4 and CD8 T cells did not change significantly in the tumor at day 8, except for mice treated with HDC aPD-1/CDN-NPs (**Fig. 5D,E**), with insignificant differences in the activation of either T cell population observed as indicated by CD69 expression (**Fig. S12**). However, for STING-containing therapies, CD8^+^ T cells were less exhausted, with a lower percentage of PD-1^+^TIM-3^+^ expression observed compared to empty hydrogel, particularly for triple combination therapy (**Fig. 5F,G**). No changes to effector memory or central memory populations were observed in the tumor, the tdLNs, or the spleen following treatments on day 8 (**Fig. S12**). The contribution of immune effector NK and T cells to therapeutic activity was further delineated using antibody-mediated NK cell and T cell depletion assays (**Fig. 5H**). CD8 T cell-depleted mice treated with triple therapy showed delayed tumor growth and significantly increased median survival relative to untreated mice but had reduced long-term survival (14.3%) and median survival (23 days) relative to mice treated with isotype antibodies and triple hydrogel therapy (undefined, 87.5%) (**Fig. 5I**). NK cell-depleted mice also showed a complete loss of therapy efficacy, with no significant median survival extension observed in mice and only one mouse surviving long-term (16.7%) (**Fig. 5J**). Together, these studies indicate that the activity of both NK cells and CD8 T cells contributes to the increased long-term survival observed in HDC triple therapy treated mice, which is consistent with what has been observed for the clearance of metastatic lesions by STING therapy^35^. These studies highlight the importance of both changes in the frequency and state of effector immune cells for HDC combination immunotherapy efficacy for the treatment of brain tumors.

## Conclusions

The development of effective anti-tumor therapies in GBM relies on the ability to overcome multiple mechanisms of intrinsic and adaptive tumor resistance. This makes treatment of GBM with monotherapies challenging and exacerbates existing limitations on drug selection made by poor systemic delivery to the brain. We reasoned that the synchronous coordination of combination therapy delivery with biological processes in the TME would allow for synergistic effects that overcome drug-resistance mechanisms without inducing autoimmunity. However, currently, there are a lack of clinically translatable drug delivery systems that can effectively control the timing and duration of combination therapies in the brain over several weeks. Here, we present injectable HDCs that engineer material-drug interactions through endogenous or engineered amines to control the intracranial delivery of combination therapy following adhesion to brain tissue, which allowed us to overcome adaptive resistance to monotherapy while preventing neurotoxic responses (**Fig. 4.2B-D**). This is distinct from previous brain delivery systems^17–19,36^ and can leverage emerging antibody-drug conjugate and NP polymer chemistries to expand the range of achievable profiles in a commercially scalable fashion. *In vivo* orthotopic studies demonstrated that HDCs tuned the delivery of multiple payloads while limiting the exposure of therapies to the contralateral hemisphere or peripheral organs (**Fig. 4.2H-K**). When used to deliver an *in situ* vaccine combination therapy over the course of several weeks, HDCs potently eliminated established brain tumors and enhanced long-term survival upon rechallenge, including in murine tumors resistant to STING agonist and ICB monotherapy, through the activation of diverse anti-tumor immune responses following a single dose (**Fig. 4.3E,J**). The prolonged hydrogel release of immunomodulatory therapies further reduced the potential for immune escape, leading to enhanced mouse survival relative to intratumoral free therapy (**Fig. 4.3E**). Mechanistically, during the first phase of release, HDC triple therapy more potently and more efficiently activated APCs and reprogrammed tumor-associated myeloid cells than did monotherapies, through the direct (STING agonist) and indirect (ICD-induced) activation of the cGAS-STING pathway in host cells (**Fig. 4.4A-C**). This enabled effective anti-tumor immunity that depended on the engagement of both MHC-dependent and MHC-independent effector cells, where reduced T cell exhaustion and sustained NK cell activation were observed as late as eight days post-therapy (**Fig. 4.5 C,F,G**). Sustained local release of aPD-1 from HDCs further prevented adaptive immune resistance by blocking the action of upregulated immune checkpoint molecules within the tumor following innate immune stimulation and subsequent production of interferons (**Fig. 4.4Q**). Collectively, our results emphasize the importance of tuning combination therapy release kinetics to align with the time scale of biological processes to produce effective, durable immune responses without inducing autoimmunity or neurotoxicity.

While our results highlight the power of efficiently converting the TME to effect local immune responses using controlled biomaterial-based delivery systems, local administration can also reduce therapy exposure to peripheral immune organs, which may impact T cell antigen priming and reactive clonotype expansion^37^. Though locoregional delivery improved ICB efficacy in mice in peripheral tumors by improving TME and tdLN exposure via lymphatic drainge^38^, lymphatic drainage and drug elimination from the CNS is distinct from the periphery^39^. Ineffective CNS drainage (of antigens) from the brain into the tdLNs impaired systemic ICB responses to primary murine brain tumors in prior studies^40^. Furthermore, the presence of peripheral lesions in the metastatic setting^41^ or priming from peripheral organs that share draining LNs demonstrated improved brain tumor responses to vaccines in mice^42^, suggesting the importance of peripheral immune T cell education. However, our results suggest minimal tdLN involvement at the timepoints studied. Other studies have pointed to the potential of local immune niches for T cell priming and activation in ICB-resistant gliomas^43^. Additional studies are thus required to dissect the interplay and contributions of local and systemic immune responses in the setting of primary brain tumors to further optimize pharmacokinetic profiles to bolster antigen-specific immune responses. Regardless, it is intriguing to consider the potential of a locally contained response given the distinct anatomical constraints of coordinating immunity across the CNS and periphery. Future work may leverage emerging knowledge about local CNS-immune compartments, such as the dural sinuses^33^ or skull bone marrow^44,45^, to realize this potential.

Collectively, the current work demonstrates the significant impact that therapy selection and biomaterial-mediated therapy release kinetics have on the evolution of effective immune responses to treat primary brain tumors. Our studies teach us that a monotherapy, and perhaps even dual therapy, is not sufficient in eliminating aggressive brain tumors such as GBM, which incite numerous immune evasion mechanisms. We show that therapies with complementary immune activation mechanisms are required over prolonged duration to enable not only immune activation but to also overcome the succeeding immune regulatory phase over a period of weeks. Our findings demonstrate the critical dependence of multiple interconnected MHC-dependent and MHC-independent immune axes on mounting and sustaining effective anti-tumor responses and emphasize how biomaterials can be used to engineer the complex interactions between innate and adaptive immune activation to generate these therapeutic responses. Future studies to assess this delivery technology and approach in large animals and in a first-in-human clinical trial are being planned. Together, these data establish our adhesive HDC platform as a promising technology for engineering therapeutic immunity to brain tumors and other therapy-resistant lesions.

## Materials and Methods

### Materials

All chemicals and cell culture materials were purchased from Sigma-Aldrich unless otherwise stated. Arginine peptide was obtained from CPC Scientific. Generation 5 PAMAM dendrimer was purchased from Dendritech. DOX hydrochloride was purchased from Cayman Chemicals. TCDN-2 and TCDN-2 maleimide were kindly provided by Takeda Pharmaceuticals. Murine anti-PD-1 (clone RMP1-14), anti-NK1.1 (clone PK136), anti-CD8α (clone 2.43), anti-CD4 (clone GK1.5), IgG2a isotype antibody (clone C1.18.4), and IgG2b isotype antibody (clone LTF-2) were purchased from BioXCell.

### Cell lines

GL261-luciferase and CT-2A-luciferase cells were cultured in Dulbecco’s minimum essential medium (DMEM) supplemented with 10% (v/v) fetal bovine serum (FBS), 100 U/mL penicillin and 100 ug/mL streptomycin. BV.2 cells were purchased from Cell Line Services and cultured in RPMI-1640 supplemented with 10% (v/v) FBS. bEND.3 cells (ATCC) were cultured in DMEM supplemented with 10% (v/v) FBS per the provider’s instructions. All cell lines were maintained in a humidified incubator at 37°C, 5% CO_2_.

### Animals

Wildtype C57BL/6 mice (6-8 weeks, strain 027) were purchased from Charles River Laboratories. C57BL/6-Tg(TcraTcrb)1100Mjb/J mice (OT-1, 6-10 weeks, strain 003831), C57BL/6J-*Sting1^gt^*/J mice (Goldenticket/STING^Gt^, 6-8 weeks, strain 017537), and B6.129S(C)-*Batf3^tm1Kmm^*/J mice (Batf3^-^, 6-8 weeks, strain 013755) were purchased from Jackson Laboratories. All studies were performed according to protocols approved by the Institutional Animal Care and Use Committee (IACUC) of MIT.

### Dextran aldehyde synthesis and fluorescent labeling

Dextran aldehyde (50% oxidized) was synthesized by periodate oxidation of dextran with sodium periodate as previously described. Briefly, dextran (10 g) was dissolved in deionized water to which sodium periodate (9.3 g) was added dropwise and stirred in the dark for four hours at room temperature. The solution was then dialyzed in deionized water in a dialysis bag (3.5 kDa MWCO) for 6 days followed by lyophilization. Proton nuclear magnetic resonance was used to confirm the chemical structure of oxidized dextran, and the degree of oxidation was quantified by the hydroxylamine hydrochloride/sodium hydroxide titration assay. For hydrogel degradation studies, dextran aldehyde was fluorescently labeled with Alexa Fluor 647 (AF647) hydrazide through the formation of a hydrazone bond, which was further reduced, following cooling, using sodium cyanoborohydride to form a stable secondary amine bond. Dextran aldehyde (10 mg) was dissolved in 1 mL of 0.1 M phosphate buffer, pH 7.5, and then the fluorescent dye (2 mg) was added dropwise to the solution and stirred for 1 hour at room temperature. The reaction solution was then cooled in an ice water bath and 1 mL of 30 mM sodium cyanoborohydride in 50 mM carbonate buffer, pH 8.5, was added and the reductive amination reaction proceeded for 4 hours. Following completion, the reaction was purified by dialysis in water at 4°C for five days using a 3 kDa MWCO dialysis cassette (ThermoFisher). The purified product was then lyophilized.

### Synthesis and characterization of pBAE CDN-NPs

CDN-NPs were synthesized as we previously described^22^. Briefly, acrylate-terminated pBAE backbone polymers (C6 and C32) were synthesized by an addition reaction of primary amines (C6: 5-amino-1-pentanol, hexylamine; C32: 5-amino-1-pentanol) to an excess of 1,4-butanediol diacrylate. Conjugated pBAE CDN polymer was generated by Diels-Alder reaction between the CDN’s maleimide group and the furan group of 2-methyl-3-furanthiol-functionalized C6 pBAE. Polypeptide-modified pBAEs were synthesized by end-modification of acrylate-terminated C32 pBAE with thiol-terminated arginine polypeptide (H-Cys-Arg-Arg-Arg-NH_2_, CPC Scientific). All structures were confirmed by ^1^H-NMR (400 MHz Varian) following dissolution in an appropriate deuterated solution. Mixtures of polypeptide- and CDN-modified pBAEs were nanoprecipitated to generate discrete NPs that were further PEGylated and purified by centrifugal filtration. The extent of CDN loading was quantified by liquid chromatography and tandem mass spectrometry (LC-MS/MS) using a triple quadrupole instrument and the size and surface charge of the CDN-NPs were characterized by dynamic light scattering (Zetasizer Nano ZS, Malvern Instruments).

### BMDM and BMDC isolation and culture

Mice (C57BL/6, 8-10 weeks old) were euthanized by carbon dioxide inhalation and their intact femurs were resected following tissue debridement and flushed with ice cold PBS under sterile conditions to obtain bone marrow progenitor cells. The resultant cell population was filtered through a 40-um cell strainer and red blood cells were lysed using ACK and centrifuged according to the manufacturer’s protocol. To generate BMDMs, 5-8×10^6^ monocytes were cultured in 25 mL of BMM+ media (DMEM/F12 media supplemented with 10% (v/v) FBS, 1% (v/v) PenStrep, 5% (v/v) Glutamax (200 mM), and 20 ng/mL recombinant murine macrophage colony stimulating factor) in a T175 non-tissue-culture-treated flask. On day 3, the flask was supplemented with 25 mL of fresh BMM+ media. On day 6, 25 mL of culture media was replaced with 25 mL of fresh BMM^+^ media. On day 8, the media was removed and BMDMs were washed, harvested using Accumax, centrifuged, and resuspended in BMM^+^ media prior to seeding for experiments. To generate BMDCs, 2×10^6^ bone marrow cells were added to non-tissue culture-treated petri dishes and cultured in 10 mL of BMG^+^ media (RPMI-1640 supplemented with 10% (v/v) FBS, 1% (v/v) penicillin/streptomycin (P/S), and recombinant murine granulocyte-macrophage colony-stimulating factor (GM-CSF, 20 ng/mL)). On day 3, the flask was supplemented with 10 mL of fresh BMG^+^ media. On days 6 and 8, 10 mL of the consumed culture media was centrifuged, and the resultant cell pellet was resuspended in 10 mL of fresh BMG+ media and added back to the petri dish. On day 10, BMDCs were collected by pipetting and gently washing the plate with cold PBS.

### NK and T cell isolation

Female OT-I mice (6-8 weeks old) were euthanized, and their spleens were extracted and washed in cold PBS under sterile conditions. The spleens were then mechanically dissociated through a cell strainer and erythrocytes were removed using ACK lysis buffer (Gibco). T cells were isolated from all splenocytes using the EasySep™ Mouse T cell isolation kit (STEMCELL technologies) following the manufacturer’s instructions. T cells were cultured in RPMI-1640 supplemented with 10% (v/v) FBS, 2 mM L-glutamine (Gibco), 1% (v/v) P/S, 20 mM HEPES, 1 mM sodium pyruvate, 1x MEM nonessential amino acids, 55 μM β-mercaptoethanol, and 10 ng/mL murine IL-2 (Peprotech). For NK cells, splenocytes were isolated from C57BL6 mice following the same process except that the EasySep™ Mouse NK cell isolation kit was used instead. Following isolation, NK cells were further expanded by culturing them in MEMa (nucleosides, no phenol red) medium (Thermo) containing 20% (v/v) FBS, 1% (v/v) Pen-Strep, 50 μM 2-mercaptoethanol, and 150 ng/mL recombinant murine IL-15 (Thermo).

### BMDM and microglia polarization

BMDM or BV.2 microglia cells were seeded in 24-well plates and allowed to incubate overnight. To obtain M2 polarized cells, the media in the wells was replaced with media supplemented with IL-13 (10 ng/mL) and IL-4 (25 ng/mL) and the cells were incubated for an additional 24 hours. The cells were then washed with PBS and treated with control media or media containing CDN-NPs for 24 hours. After treatment, cells were detached with Accumax, harvested, and stained for flow cytometry analysis.

### BMDM, BMDC, and microglia activation

Bone-marrow derived cells or microglia were seeded in 24-well plates at 200,000 cells per well then treated with CDN-NPs or PBS. After 24 h, the cells were harvested, washed, blocked with anti-CD16/CD32 (BioLegend), stained, fixed, and analyzed by flow cytometry. For combination co-culture activation studies, tumor cells were first seeded and treated with DOX or PBS for 48 h. Following incubation tumor cells were washed and co-cultured with bone-marrow derived cells. Co-cultures were treated with aPD-1 and/or CDN-NPs at indicated concentrations for 24 h. Following incubation, cells were harvested, washed, blocked with anti-CD16/CD32 (BioLegend), stained, fixed, and analyzed by flow cytometry. Cells were stained with anti-CD45-BV785, anti-CD11c-BV421, anti-F4/80-BUV395, anti-CD11b-BV421, anti-MHCII-BV605, anti-CD206 PE, anti-CD80-APC, anti-CD86-FITC, and Fixable Live/Dead.

### Phagocytosis

Tumor cells were labeled with CellTrace™ Far Red proliferation dye (ThermoFisher), then plated and allowed to adhere overnight. Tumor cells were treated with the indicated therapies for 48 hours at 37°C. Bone-marrow derived macrophages or dendritic cells were then co-cultured with tumor cells (following treatment and PBS wash) for 4 hours at 37°C. Following incubation, cells were collected on ice, washed with 1x HBSS, and treated with an FcX blocking antibody. Cells were then stained with BV421 CD11b (BMDMs) or CD11c (BMDC) antibody and analyzed by flow cytometry where phagocytosis was reported as the percentage of APC^+^ cells within the BV421 population.

### Antigen cross presentation

OVA-expressing tumor cells were seeded in well plates and allowed to adhere overnight prior to treatment with DOX for 48 hours. Following incubation, tumor cells were washed, co-cultured with BMDCs and treated with immunotherapies (CDN-NPs and/or aPD-1) for 24 h. Cells were then harvested using Accumax, washed twice in PBS, and resuspended in cell staining buffer (BioLegend). Cell suspensions were treated with FcX blocking antibody, stained with APC-labeled anti-SIINFEKL/H-2Kb, FITC-anti-CD11c, and fixable violet live-dead stain, and fixed. Antigen presentation was analyzed by flow cytometry (Symphony, BD) and was reported as the percentage of APC^+^ cells within live FITC^+^ BMDCs.

### T cell proliferation

Tumor cells were plated, treated, and co-cultured with BMDCs as previously described in the antigen cross presentation assay. Naïve CD8 T cells were isolated from WT or OT-I mice as described above and stained with CellTrace™ Far Red proliferation dye (for proliferation studies only) prior to being added to tumor-BMDC co-cultures for 48-72 h. Cells were then harvested and stained for flow cytometry analysis. For both analyses, cells were stained with anti-CD3 and anti-CD8. Proliferation was reported as the percent of diluted CD8 T cells relative to total CD8 T cells (CD3^+^CD8^+^). Maturation markers were used to distinguish naïve (CD44^low^CD62L^high^), central memory (CD44^high^CD62L^high^), and effector memory (CD44^high^CD62L^low^) populations within the overall CD8 T cell population (CD3^+^CD8^+^).

### Establishment of orthotopic murine GBM tumors in vivo

Syngeneic orthotopic mouse models of glioblastoma were established by implanting 1.3×10^6^ GL261 or CT-2A luciferase-expressing GBM cells in the left hemisphere of C57BL6 mice (6-8 weeks old, Charles River) using a stereotactic device (Stoelting). Briefly, mice were administered sustained release buprenorphine analgesia (1 mg/kg). Mice were then anesthetized with inhalable isoflurane, shaved to remove all hair from their scalps, and immobilized on a stereotactic frame. Following multiple cycles of betadine/isopropyl alcohol scrub, a small incision was made to expose the skull, and a 1-mm burr hole was drilled at two millimeters lateral and two millimeters posterior to the bregma. Cells were injected two millimeters deep from the outer border of the cranium using a 10 uL microsyringe (Hamilton Company) at a speed of 1 uL/min. The needle remained in place for 1-2 minutes following injection and was removed slowly. The incision was closed with sutures and the mice were kept on warming pad and followed through complete recovery of ambulation. On the day prior to treatment (Day 6 for CT-2A or Day 9 for GL261), mice were evaluated for tumor burden using bioluminescence imaging by IVIS following intraperitoneal administration of D-luciferin (150 mg/kg in DPBS, Perkin Elmer).

### In vivo therapeutic efficacy studies

Following confirmation of tumor establishment, mice were randomly stratified into treatment or control groups. Mice were prepared for surgery as described above. The incision was reopened and the burr hole from tumor implantation was identified and redrilled. Hydrogels containing therapies or PBS (empty) or free drug solutions were injected intratumorally using a stereotactic instrument on Day 7 (CT-2A) or Day 10 (GL261). Therapeutic doses were DOX (15 ug), CDN-NPs (75 ng CDN), and/or aPD-1 (20 ug) as identified by maximum dose tolerability studies (**Fig. S6**). Mice were anesthetized with isoflurane via inhalation throughout the duration of the procedure and were given extended-release buprenorphine (1 mg/kg) for analgesia. Mice were tracked for body weight (>20% weight loss), body condition (score < 2), development of neurological symptoms, and tumor growth by bioluminescence imaging using an IVIS. For T cell depletion studies, anti-CD8α (clone 2.43, BioXCell) or anti-CD4 (clone GK1.5, BioXCell) were administered intraperitoneally at a dose of 400 ug/dose, three times per week for the duration of the study starting 48 h before treatment. For NK cell depletion studies, anti-NK1.1 (clone PK136, BioXCell) was administered intraperitoneally at a dose of 400 ug every four days, starting 48 h before treatment.

### In vivo cytokine quantification

Mice were implanted with tumors as described in the model establishment section. Tumor growth in mice was confirmed and treated as described in the in vivo efficacy studies. At the specified time point, blood was collected into heparinized tubes from mice by submandibular cheek bleeds. The collected blood was centrifuged at 1,000 x *g* for 20 minutes at 4°C and the plasma was collected for analysis. Following blood collection, mice were administered a lethal anesthetic dose of ketamine/xylazine cocktail and were completely perfused with heparanized PBS by cardiac puncture, using the color of the liver and of the fluid exiting the mouse to indicate the extent of the perfusion. The tumor, tdLNs, and spleens of the mice were then extracted and weighed. The tissues were homogenized in lysis buffer (T-PER Tissue Protein Extraction Reagent, Thermo Scientific) containing 1% Halt protease and phosphatase inhibitor cocktail using a Precellys Tissue Homogenizer (Bertin Technologies). Lysates were centrifuged at 10,000 x *g* for 10 minutes and the supernatant was collected for cytokine concentration analysis by the LEGENDplex™ mouse inflammation panel (13-plex, Biolegend) following the manufacturer’s instruction. The total protein concentrations in all tissue supernatants were quantified using the micro bicinchoninic acid assay (Thermofisher Scientific).

### In vivo biodistribution and cell uptake studies

Hydrogels were loaded with either triple therapy containing fluorescently-labeled CDN-NPs or with PBS and injected into C57BL/6 mice bearing CT-2A or GL261 tumors as described above. At specified timepoints, mice were perfused with heparanized PBS and their brain, tdLNs, and major organs (liver, lungs, heart, kidneys, spleen) were collected on ice and imaged for fluorescence using an IVIS. To determine the cellular populations that internalized CDN-NPs in the tumor, mice were perfused and sacrificed 24 hours after hydrogel treatment and their brains were collected and dissociated into single cell suspensions. Cells were then blocked with FcX antibody, stained and fixed prior to resuspension in cell staining buffer (BioLegend) for spectral flow cytometry analysis using a Sony ID7000 instrument. Data was spectrally unmixed and processed using the Sony ID7000 analysis software and analyzed in FlowJo (v10.8.1). For assessment of the release of DOX from the hydrogel *in vivo*, mice were implanted with tumors and treated as per the CT-2A model then sacrificed on predetermined days and their brains and plasma were harvested and processed for LC/MS-MS analysis.

### Immunophenotyping Flow Cytometry

Mice were implanted with tumors, assessed for tumor burden, and treated as previously described. After the specified time, brain tumors, tdLNs, and spleens were collected as described above. For brain tumor samples, following extraction, the tissues were minced in enzymatic digestion buffer (RPMI containing 1 mg/mL collagenase D and 0.25 mg/mL DNaseI) and then incubated for 30 minutes at 37°C with intermittent pipetting. The samples were then triturated through 40 um cell strainers with cold HBSS and centrifuged. Myelin and other processing debris was removed using density gradient centrifugation with a debris removal solution (Miltenyi) following the manufacturer’s instructions to yield the final single cell suspension for counting and further processing. The spleen and tdLNs were processed into single cell suspensions by mechanical dissociation and filtered through a 40 um cell strainer. The spleen cell suspensions were further treated with ACK lysis buffer to remove erythrocytes. The resulting cell pellets were resuspended in cell staining buffer (BioLegend), filtered, and counted. Cell suspensions were plated at 2 x 10^6^ cells per well, blocked with anti-CD16/CD32 antibody (BioLegend), stained and fixed. For intracellular stains, cells were permeabilized and fixed after staining using a cytofix/cytoperm kit (BD) following the manufacturer’s instructions. Cells were stained with a combination of the following antibodies: anti-CD45-AlexaFluor700 (Biolegend), anti-CD3-BUV395 (BD), anti-CD4-BUV496 (BD), anti-CD8-BUV737 (BD), anti-FOXP3-PE (BD), anti-TIM3-BV421 (BioLegend), anti-PD1-FITC (BioLegend), anti-CD69-BV605 (BioLegend), anti-CD44-BV786 (BioLegend), anti-CD62L-PE-Cy7 (BD), anti-NK1.1-APC (BioLegend), Fixable Live/Dead Near IR (Invivogen), anti-CD45.2-BV786 (BioLegend), anti-CD11c-APC (BioLegend), anti-MHCII-BV605 (BioLegend), anti-SiglecH-PE (BioLegend), anti-CD8α−BUV737 (BioLegend), anti-CD103-BUV395 (BD), anti-CD86-BV510 (BioLegend), anti-CD80-BUV737 (BD), anti-F4/80-APC (BioLegend), anti-CD11b-BUV395 (BD), anti-Ly6G-PerCP-Cy5.5 (BioLegend), anti-Ly6C-FITC (BioLegend), anti-CD274-BV421 (BioLegend), anti-CD206-PE (BioLegend), anti-NK1.1-BV605 (BioLegend), anti-CD11c-PE-Cy7 (BioLegend), anti-CD19-BV421 (BioLegend), anti-CD8α-BV510, anti-MHCII-BV650 (BioLegend), and anti-TCRβ-BV785 (BioLegend). All samples were run on a Sony ID7000 spectral flow cytometer and processed for autofluorescence subtraction and spectral unmixing using the instrument’s software. For T cell data, multiple samples from the same mouse were concatenated together for analysis. All flow cytometry data was analyzed using FlowJo software (v10.8.1) and representative images of the flow gating strategies for all experiments have been provided (**Fig. S4.13-15**).

### Immunohistochemistry

Mice were implanted with tumors, assessed for tumor burden, and treated as previously described. At select timepoints, mice were administered a lethal anesthetic dose of ketamine/xylazine cocktail and transcardially perfused with heparanized saline. The brains of mice were resected, rinsed in sterile PBS, and fixed in formalin overnight. The brains were then precipitated in 15% (w/v) sucrose for 6-12 hours, then in 30% (w/v) sucrose overnight prior to embedding in OCT (Tissue-Tek, Sakura) and freezing at -80°C. Frozen tissues were sectioned using a cryostat instrument. Several sections from each tissue were stained using DAPI, mounted on poly-D-lysine coated slides, cover-slipped, and dried overnight prior to imaging using confocal microscopy. For tolerability studies, the major organs, including brain, lungs, heart, liver, spleen, and kidneys, were extracted, and fixed in formalin for 24 hours. Samples were then dehydrated, cleared with xylenes, and embedded in paraffin prior to sectioning using a microtome. Sections were stained with hematoxylin & eosin and evaluated using light microscopy by a trained pathologist who was blinded to study group treatments.

### Statistical Analyses

Statistical analyses were performed using GraphPad Prism 10 (GraphPad Software). All data are reported as the mean ± the standard error of the mean (s.e.m.) unless otherwise specified. For *in vitro* experiments at least three biological replicates were used per condition per experiment. Statistical significance was determined as follows: a parametric two-tailed t test was performed for the comparison of two groups at single timepoint; a one-way analysis of variance (ANOVA) was performed for the comparison of three or more groups; and a two-way ANOVA was performed when there were three or more groups and multiple timepoints. For in vivo studies, the Kruskal-Wallis test with uncorrected Dunn’s test was used for multiple comparisons between groups. Kaplan-Meier survival curves were generated and compared using the log-rank Mantel-Cox test.

## Author Contributions

MZD and NA conceived the study. MZD designed the experiments. MZD, DD, AL, IL, SC, and AH performed the *in vitro* experiments. MZD performed the *in vivo* surgeries, and subsequent analyses with assistance from AMC, DD, PD, MP, NP, and AL. BT performed the MTD and hydrogel degradation in vivo experiments. BT and HB provided input on neurosurgical experiments. MZD and NA drafted and finalized the manuscript with input from all authors.

## Acknowledgements

The authors thank the Robert A. Swanson (1969) Biotechnology Center at the Koch Institute for Integrative Cancer Research at the Massachusetts Institute of Technology (NCI Cancer Center Support Grant P30-CA014051) for assistance with animal experiments and facilities, particularly the B76 animal facility, flow cytometry, histology, and nanomaterials characterization core staff. We thank A. Hayward for assistance with surgical model training, D. Markus and A. Lytton-Jean for cryo-SEM assistance, J. Wyckoff for confocal imaging training, K. Cormier for assistance with histology, and G. Pardis for assistance with flow panel design. We thank E. Zigon at the Wyss Institute for assistance with spectral flow cytometry. Biorender was used to create several schematics.

## Funding

M.Z.D. acknowledges a National Science Foundation Graduate Research Fellowship and fellowships from the MIT School of Engineering (Evergreen Graduate Innovation Fellowship) and MIT HST (Price Innovation Fellowship). This project was supported in part by funding from the MIT Deshpande Center for Technological Innovation and the Koch Institute Cancer Center Support Grant P30-CA14051 from the National Cancer Institute. Any opinion, findings, and conclusions or recommendations expressed in this material are those of the authors and do not necessarily reflect the views of the National Science Foundation.

## Notes

### Competing Interest Statement

The authors have declared no competing interest.

